# Protein enhanced NIR-IIb emission of indocyanine green for functional bioimaging

**DOI:** 10.1101/2020.05.30.125104

**Authors:** Mubin He, Di Wu, Yuhuang Zhang, Xiaoxiao Fan, Hui Lin, Jun Qian

**Author notes:** Corresponding author: Jun Qian.

## Abstract

Fluorescence imaging performed in the 1500-1700 nm spectral range (labeled as near-infrared IIb, NIR-IIb) promises high imaging contrast and spatial resolution for its little photon scattering effect and minimum auto-fluorescence. Though inorganic and organic probes have been developed for NIR-IIb bioimaging, most are in preclinical stage, hampering further clinical application. Herein, we showed that indocyanine green (ICG), an US Food and Drug Administration (FDA)-approved agent, exhibited remarkable amount of NIR-IIb emission when dissolved into different protein solutions, including human serum albumin, rat bile, and fetal bovine serum. We performed fluorescence imaging in NIR-IIb window to visualize structures of lymph system, extrahepatic biliary tract and cerebrovascular. Results demonstrated that proteins promoted NIR-IIb emission of ICG *in vivo* and that NIR-IIb imaging with ICG preserved higher signal-to-background ratio (SBR) and spatial resolution compared with the conventional near-infrared II (NIR-II) fluorescence imaging. Our findings confirm that NIR-IIb fluorescence imaging can be successfully performed using the clinically approved agent ICG. Further clinical application in NIR-IIb region would hopefully be carried out with appropriate ICG-protein solutions.

## 1. Introduction

Fluorescence imaging in the near-infrared window (NIR:760-2400 nm) exhibits prominent advantages including real-time imaging, free radiation and superb spatial resolution, compared with traditional imaging techniques such as computed X-ray tomography (CT), positron emission tomography (PET) and magnetic imaging (MRI) ^[1-5]^. Among NIR spectral region, imaging performed in the second NIR window (NIR-II, 900-1700 nm) outperformed that in the first near-infrared window (NIR-I, 760-900 nm) as the photon scattering effects and endogenous tissue auto-fluorescence signal could be significantly reduced at longer wavelength region ^[5-8]^. Moreover, spectral region of 1500-1700 nm (NIR-IIb) promises even larger tissue penetrating depth and minimum auto-fluorescence due to its longest wavelength in NIR-II window ^[9]^. Diversified emitters have been developed for visualizing anatomical structures in recent years ^[10]^. However, inorganic nanomaterials including rare-earth doped nanoparticles ^[11, 12]^ and quantum dots ^[13]^ confronted long-term cytotoxicity troubles. Though recently developed organic fluorophores ^[14-17]^ exhibited emission beyond 1500 nm, they also encountered low molar extinction coefficients, or long clinical approval process.

Indocyanine green (ICG), approved by the US Food and Drug Administration (FDA), was a conventional NIR-I dye, which had been widely applied for clinic ^[18]^. ICG was reported to emit NIR-II fluorescence suitable for noninvasive imaging in blood and lymph vessels in mice ^[19]^. Recent clinical applications based on NIR-II fluorescence of ICG have demonstrated the brilliant sensitivity in detecting the primary tumor and extrahepatic metastases ^[20]^. Since the inherent defects including poor photostability ^[6, 21]^ in water, rapid plasma clearance rate in circulation and strong background signal due to its accumulation in liver have impeded ICG’s further clinical application, efforts have been tried to stabilize the fluorescent agent. Liposomal formation of ICG and incorporating ICG into PEG have been reported to increase ICG’s half-life period in blood ^[19, 22]^. ICG in fetal bovine serum (FBS) also demonstrated enhanced NIR-II fluorescence intensity ^[23]^. So far, ICG fluorescence-guided surgeries are still mainly performed in the NIR-I and NIR-II (specifically in 1000-1400 nm) spectral region.

Here, the fluorescence intensity of ICG dilutions in human serum albumin (HSA), rat bile (containing mucin and plasma proteins) and FBS in the NIR-IIb region were measured. NIR-IIb fluorescence imaging of lymph nodes, extrahepatic biliary tract and cerebrovascular structures with ICG *in vivo* were also conducted and compared with those in the NIR-II window. The primary objective of the current study was to evaluate the feasibility of performing fluorescence imaging in the NIR-IIb window using a clinically approved agent.

## 2. Materials and Methods

### 2.1. Materials

Indocyanine green (ICG) was acquired from Sir Run-Run Shaw Hospital, School of Medicine, Zhejiang University. Fetal bovine serum (FBS) and human serum albumin (HSA) were purchased from Dalian Meilun Biotechnology Co., LTD. Phosphate-buffered saline (PBS) was attained from Sinopharm Chemical Reagent Co., LTD., China. Rat Bile *in vitro* was extracted from live rat (∼8 weeks old). Deionized (DI) water was extracted from an Eco-Q15 deionized water system (Shanghai Hitech Instruments Co., Ltd.).

### 2.2. Absorption and photoluminescence spectra characterization

The absorption spectra of ICG dilutions in DI water, 4% HSA, rat bile and FBS were measured by Shimadzu UV-2550 UV-vis-NIR scanning spectrophotometer. The photoluminescence spectra were recorded by a NIR2000C spectrometer (Everuping Optics Corporation).

### 2.3. Optical system for NIR-IIb fluorescence macro-imaging

The NIR-IIb fluorescence macro-imaging optical system contained excitation part and signal collection light part (Fig.S1). In the excitation part, a 793 nm laser with collimator was stained on an optical pole. The collimator had an 850 short-pass (850 SP Thorlabs) filter which cut off the longer wavelength of light source. In the signal collection part, an electronic-cooling 2D (640 pixels × 512 pixels) InGaAs camera (TEKWIN SYSTEM, China) which has the good sensitivity at the spectral range from 900 nm to 1700 nm, was immobilized in an aluminum frame. Below the camera was a prime lens (focal length: 35 mm, Edmund Optics) with antireflection (AR) coating at 800-2000 nm. InGaAs camera thus only collected NIR-IIb (1500-1700 nm) fluorescence signals when equipped with a 1500 nm LP filter (Thorlabs). The distance between the camera and imaging plane could be adjusted in order to focus.

### 2.4. Optical system for NIR-IIb fluorescence mesoscopic/microscopic imaging

The NIR-IIb fluorescence microscopic imaging optical system (NIRII-MS, Sunnyoptical) adds 900 nm long-pass dichroic mirror (DMLP) and objectives compared to macroscopic imaging system (Fig.S2). The tube lens had 1000-1700 nm anti-reflection. A 793 nm laser was equipped as excitation source. After 793 nm laser beams reflect from a 900 nm long-pass dichroic mirror (DMLP) and pass through an objective, the observed sample is illuminated. The objective could be an air objective lens (LSM03, WD= 25.1 mm, Thorlabs or LMS05, WD=36 mm, Thorlabs), or an infrared transmission water-immersed object (XLPLN25XWMP2, 25×, NA=1.05, Olympus). NIR-IIb fluorescence images of the sample were recorded with the InGaAs camera after passing through a 1500 nm long-pass filter (Thorlabs).

### 2.5. Ex vivo fluorescence and photostability test

ICG (0.1 mg/mL) was dissolved in DI water, 4% HSA, rat bile and FBS respectively. Four centrifuge tubes which respectively contained 100 µL above-mentioned solution, were irradiated by a 793 nm CW laser with 60 mW/cm^2^ for 200 min uninterruptedly. The fluorescence images in NIR-IIb region were recorded every 5 min by the InGaAs camera (exposure time= 100 ms) under NIR-IIb fluorescence macro-imaging optical system.

### 2.6. Sacral/popliteal lymph node NIR-IIb imaging

Sacral/popliteal lymph node NIR-IIb imaging was also performed by the previous NIR-IIb fluorescence macro-imaging system. ICG (2 mg/mL, 50 μL) in DI water, 2% HSA, and 4% HSA were injected into each digit and the footpad of anesthetic BALB/c nude mice (4∼6 weeks female) respectively. Mice were then placed on the imaging platform and taped safely in a position where the Sacral/ popliteal lymph node were observed clearly on InGaAs camera. To attain 1000 nm LP and 1300 nm LP macro-image, the filter in front of the camera test surface were changed from 1500 nm LP to 1000 nm LP and 1300 nm LP accordingly. Imaging conditions: 793 nm laser power= 22 mW/cm^2^ (1000 nm LP), 22 mW/cm^2^ (1300 nm LP), and 28 mW/cm^2^ (1500 nm LP); exposure time= 1 ms (1000 nm LP), 200 ms (1300 nm LP), 400 ms (1500 nm LP).

### 2.7. In vivo NIR-IIb fluorescence biliary imaging

*In vivo* NIR-IIb fluorescence biliary imaging was fulfilled with previous NIR-IIb fluorescence macro-imaging system and NIR-IIb fluorescence mesoscopic imaging optical system. The mouse (6∼7 weeks female ICR) was anesthetized via intraperitoneal injection. ICG (0.2 mg/mL, 200 µL) was injected through caudal vein. Biliary tract was imaged via the optical system after laparotomy was carried out. Imaging conditions: 793 nm laser power = 70 mW/cm^2^ (macroscopy), 190 mW/cm^2^ (mesoscopy 2X), 900 mW/cm^2^ (mesoscopy 4X), and 800 mW/cm^2^ (mesoscopy 5X); exposure time= 100 ms for macroscopy, mesoscopy 2X, mesoscopy 4X and mesoscopy 5X imaging.

### 2.8. In vivo NIR-IIb fluorescence cerebrovascular imaging

In vivo NIR-IIb fluorescence cerebrovascular imaging was achieved through NIR-IIb fluorescence microscopic imaging optical system. The mice (4∼6 weeks female BALB/c, divided into two groups) were performed microsurgery to remove the skin covering on the skull ^[24]^ for the intact skull imaging (group one) or the skull ^[25]^ for imaging with a cranial window (group two). The NIR-IIb fluorescence images were recorded immediately after ICG (2.5 mg/mL, 100 µL) was injected through caudal vein. Imaging conditions for through skull imaging with mesoscopy 5X objective: 793 nm laser power = 680 mW/cm^2^ (1000 nm LP), 1.2 W/cm^2^ (1300 nm LP), and 2.7 W/cm^2^ (1500 nm LP); exposure time = 7 ms (1000 nm LP), 300 ms (1300 nm LP), and 900 ms (1500 nm LP).

### 2.9. In vivo PTI damage observation with NIR-IIb fluorescence imaging

The mice (4∼6 weeks female BALB/c) with a cranial window were used for the PTI mouse model. At first, a micro region of the brain was selected and imaged under NIR-IIb fluorescence microscopic imaging system. After that, 40 μL of photosensitizer Rose Bengal (10 mg/mL) in PBS was infused via tail vein. Then, the same area on a capillary in the micro region was selected and illuminated for 3 min with the 532 nm laser beam (average power after the objective is 60 mW) to induce PTI damage. Finally, the micro region with PTI damage was imaged under NIR-IIb fluorescence microscopic imaging system again. Imaging conditions for 25X NIR-IIb fluorescence microscopic imaging: 793 nm laser power= 400 mW, exposure time=100 ms.

### 2.10. Ethical approval

All animal experiments performed in studies were conducted strictly in accordance with the ethical standards of the Institutional Ethical Committee of Animal Experimentation of Zhejiang University. All procedures performed in studies involving human participants were in accordance with the ethical standards of the Clinical Research Ethics Committee of Sir Run-Run Shaw Hospital of Zhejiang University.

### 2.11. Data analysis

Image J software (Version 1.6.0, National Institutes of Health, USA) was applied for quantitative analysis of each fluorescent image. Origin Pro software (Version 9.0, OriginLab Company, USA) and Adobe Illustrator CC (Version 2018) was applied for graphs generation.

## 3. Results and Discussion

### 3.1. Optical characterization

The chemical structure of ICG was presented in Fig.1a and its molar mass is 774.96 g/mol. It was reported that the emission spectrum of ICG in serum could extend to 1300 nm ^[23]^, and the tail emission spectrum of ICG in water could also prolong to NIR-IIb region at *in vitro* level ^[19]^. Nevertheless, NIR-IIb imaging of ICG *in vivo* hasn’t been carried out. We investigated the NIR-IIb fluorescence properties of ICG. The NIR-IIb fluorescence intensity measurement and photostability test were performed. Intensified NIR-IIb fluorescence signal occurred when ICG was dissolved in HSA, rat bile and FBS (Fig.1b). Quantitative analysis demonstrated that ICG in rat bile emitted 6.5-fold higher NIR-IIb fluorescence intensity than ICG in DI water. Fluorescence intensity of ICG in both FBS solution (as purchased) and 4% HSA solution (concentration = 40 mg/mL) increased to ∼ 5.2-fold compared with that in DI water (Fig.1c). Although FBS and HSA have been reported to enhance fluorescence intensity of other small-molecular NIR-II probes ^[21, 26, 27]^, the intensification of NIR-IIb fluorescence of ICG was firstly demonstrated.

**Fig. 1.**
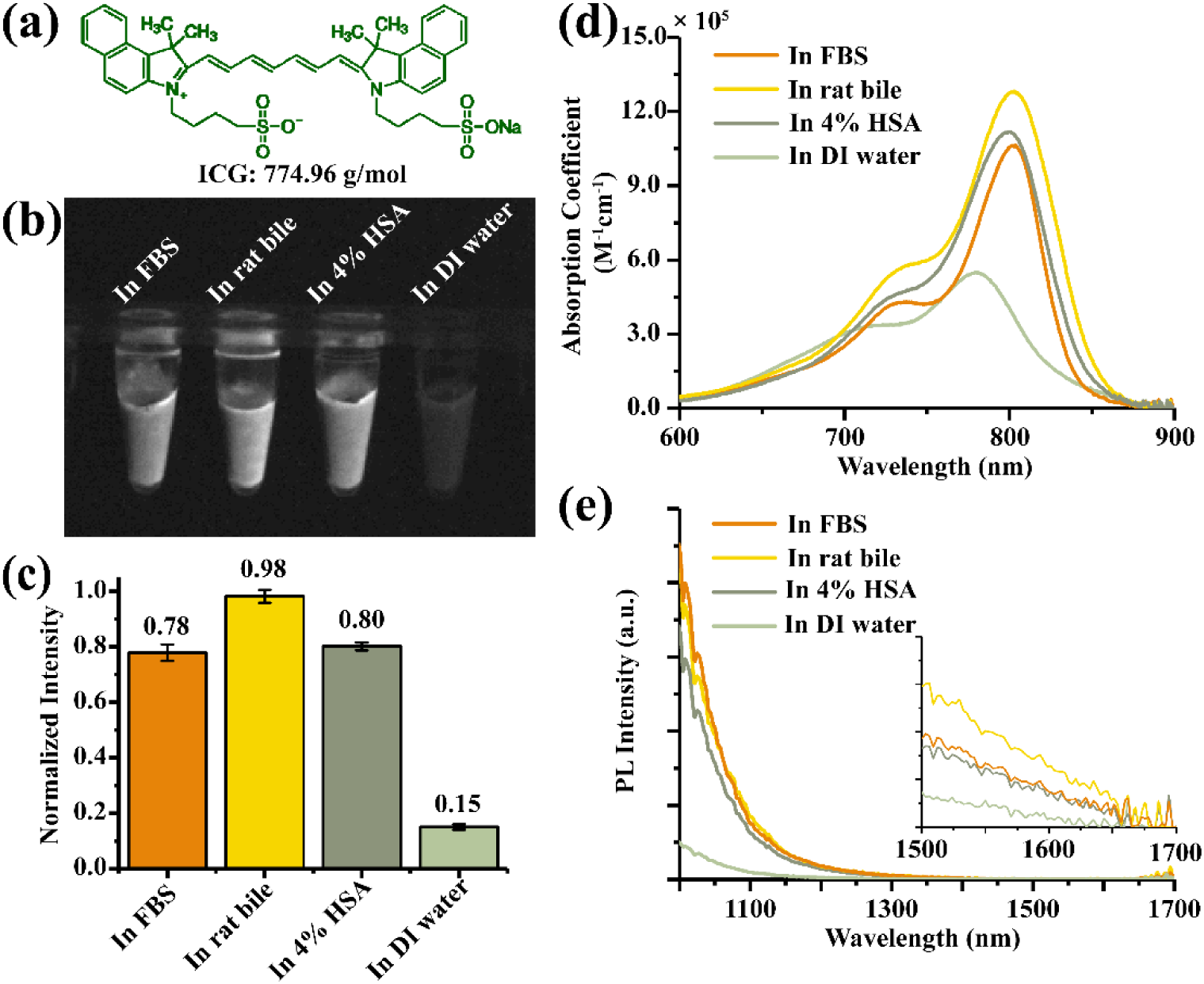
Optical characterizations of indocyanine green (ICG). (a) Molecular structure of ICG. (b) The NIR-IIb fluorescence signals of ICG (0.1 mg/mL) in FBS solution, rat bile, 4% HSA solution and DI water, under 793 nm laser excitation (60 mW/cm^2^). 1500 nm long-pass filter was adopted and the exposure time was 100 ms. (c) The calculated NIR-IIb intensities of ICG in various protein solutions, according to the images of Fig.1B. (d) The absorption coefficient of ICG in FBS solution, rat bile, 4% HSA solution and DI water. (e) The photoluminescence spectra of ICG (0.1 mg/mL) in NIR-II region; insert: spectra of ICG in NIR-IIb region. ICG (0.1 mg/mL) in DI water, 4% HSA, rat bile and FBS were excited under the same condition (793 nm laser).

ICG showed red-shifted and enhanced absorption spectra when dissolved in 4% HSA, rat bile and FBS (Fig.1d). The absorption peaked at 803.5 nm (in FBS solution), 803.0 nm (in rat bile solution) and 800.5 nm (in 4% HSA solution), red-shifting about 20 nm compared with the peak in DI water (778.5 nm). Peak absorption cross-sections of ICG in 4% HSA, rat bile and FBS increased up to ∼2.0-fold compared with that in DI water. As a result, the photoluminescence spectra of ICG in NIR-II window were also intensified under the 793 nm excitation, when dissolved in 4% HSA, rat bile and FBS (Fig.1e). The inserted spectra further confirmed the more bright fluorescence of ICG in 4% HSA, rat bile and FBS. In addition, under the continuous 793 nm laser excitation, the NIR-IIb fluorescence decay times of ICG in 4% HSA (70 min), rat bile (more than 200 min) and FBS (158 min) was longer than that in DI water (63 min) (Fig.S3). Combining the aforementioned results, we know that ICG in 4% HSA, rat bile and FBS emitted much higher and more lasting NIR-IIb fluorescence intensity than ICG in DI water, which are suitable for the following NIR-IIb fluorescence functional bioimaging.

### 3.2. NIR-IIb fluorescence imaging for lymph system

ICG has been applied for the resection of sentinel lymph nodes (SLN) during fluorescence-guided surgeries ^[28, 29]^. However, the deficiencies including poor contrast and low resolution in the NIR-I window were evident ^[21, 30]^. Organic small molecules such as CH1055 ^[30]^ and CH-4T ^[21]^ were developed in succession and allowed high SBR and resolution in NIR-II fluorescence imaging of lymph. Though it was reported NIR-II imaging of lymph vessels with ICG in water preserved contrast up to about 1400 nm, ^[19]^ imaging of lymph system with ICG in NIR-IIb region (1500-1700 nm) has not been demonstrated.

We studied the effectiveness of NIR-IIb fluorescence imaging with ICG in lymphatic drainage *in vivo*. ICG mixed with HSA (2 mg/mL, 50 μL) was injected intradermally at the rear paw of nude mice. It was evident that NIR-II (1000 nm, 1300 nm and 1500 nm long pass filter) fluorescence imaging for both lymph vessels and lymph nodes (LNs) exhibited enhanced intensity as the proportion of HSA increased from 0% to 4% (Fig.2a). Quantitative analysis of the NIR-IIb fluorescence of sciatic LNs stained with ICG in 4% HSA showed 11.3 (left) to 5.47 (right) fold increase in intensity, compared with those stained with ICG in 0% HSA (Fig.2b). Moreover, the results implied that HSA promoted ICG to flow from afferent lymph vessels to popliteal LN and efferent lymphatic vessels to sciatic LN. Thus, fluorescence imaging using ICG in 4% HSA (HSA 4% row in Fig.2a) was applied for further analysis. The lymphatic drainage of the same mouse was imaged under the 1000 nm LP, 1300 nm LP and 1500 nm LP respectively. The full width of half-maximum (FWHM) of sciatic LN beyond 1500 nm was measured as 0.842 mm, lower than those beyond 1300 nm (1.185 mm) and 1000 nm (1.346 mm) (Fig.2c). The SBR of the same LN beyond 1500 nm was calculated as 24.1, much higher than those beyond 1300 nm (15.9) and 1000 nm (7.30). Similarly, the imaging of afferent lymphatic vessels beyond 1500 nm presented highest contrast (SBR= 3.40) and resolution (FWHM= 0.530 mm) compared with those beyond 1300 nm (SBR= 2.35, FWHM= 1.057 mm) and 1000 nm (SBR= 1.32, FWHM= 0.970 mm) (Fig.2d).

**Fig. 2.**
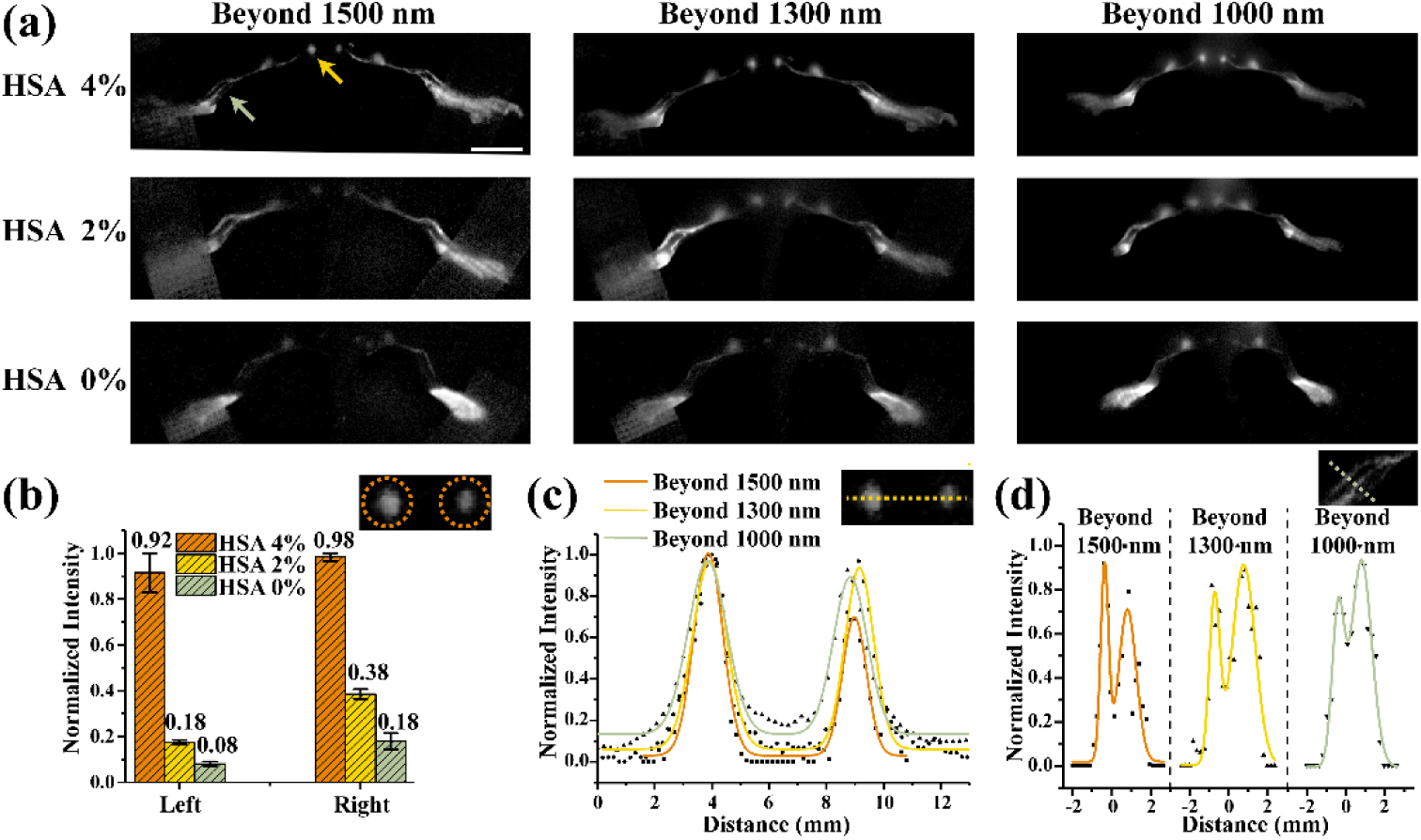
*In vivo* NIR-II fluorescence imaging of lymphatic drainage. (a) Fluorescence images of lymphatic drainage in various spectral regions. The parameters (in supplementary method) were adapted to obtain optimal image contrast. Scale bar= 10 mm. (b) NIR-IIb fluorescence intensities of sciatic lymph nodes of the mouse, which was injected with ICG in HSA solution (the proportion of HSA increased from 0% to 4%). (c) Gaussian fitting curves of the NIR-IIb fluorescence intensities of sciatic lymph nodes. The left sciatic lymph node’s FWHM= 0.842 mm (1500 nm LP), 1.185 mm (1300 nm LP), 1.346 mm (1000 nm LP); SBR= 24.1 (1500 nm LP), 15.9 (1300 nm LP), 7.30 (1000 nm LP). (d) Gaussian fitting curves of the NIR-IIb fluorescence intensities of efferent lymph vessels in left hindlimb. The left efferent lymph vessel’s FWHM= 0.530 mm (1500 nm LP), 1.057 mm (1300 nm LP), 0.970 mm (1000 nm LP); SBR= 3.40 (1500 nm LP), 2.35 (1300 nm LP), 1.32 (1000 nm LP).

Therefore, NIR-IIb fluorescence imaging of lymph drainage system assisted with ICG in HSA solution demonstrated strong potential for future surgical application.

### 3.3. NIR-IIb imaging for biliary tract

Traditional laparoscopic cholecystectomy (LC) have encountered many problems such as risks of biliary anatomical variations and bile duct injuries ^[31-33]^. NIR-I fluorescence imaging with ICG was reported in detecting bile excretion during hepatoportoenterostomy for the treatments of biliary atresia (BA) ^[34]^. A systematic review illustrated that extrahepatic bile duct visualization assisted with ICG NIR-I fluorescence system was superior to intraoperative cholangiogram ^[35]^. However, it was mentioned that high fluorescence signal of liver parenchyma have impeded the contrast of bile duct ^[18]^, making it difficult to assess the bile leakage ^[36, 37]^. Therefore, it is critically important to develop an imaging technology to visualize the biliary tract in high contrast, especially for the structures adjacent to the liver tissue.

Our previous study has confirmed the extrahepatic cholangiography in NIR-II window outperformed the conventional NIR-I imaging ^[38]^. Herein, we further investigated the capability of ICG for NIR-IIb fluorescence imaging of biliary tract at both macroscopic and mesoscopic level (Fig.3a-d). Macroscopic imaging conditions which we applied were mild: ICG dosage was 0.2 mg/mL, 200 μL; 793 nm laser excitation power was 70 mW/cm^2^; and exposure time of InGaAs camera was 100 ms. Macroscopic imaging showed that the common bile duct (CBD) was bright with high contrast compared with the liver parenchyma and duodenum (Fig.3a). Though ICG was accumulated in liver parenchyma, the fluorescence intensity of liver in NIR-IIb window was lower than that in biliary tract. We speculated that the large amount of water in liver absorbed and weakened its NIR-IIb fluorescence intensity. Additionally, cystic duct was clearly visible (FWHM= 0.299 mm and SBR= 7.56, Fig.3e). In order to precisely visualize the structure of biliary tract, three kinds of objectives with different magnifications were employed for mesoscopic imaging. The sharpness of cystic duct was apparent in Fig.3b-d. FWHM of cystic duct improved compared with macroscopic imaging. Moreover, we were able to visualize the ICG flux in the biliary tract at a speed of 25 frames per second, even there existed the interference from the rapid heart-beat (Movie S1).

**Fig. 3.**
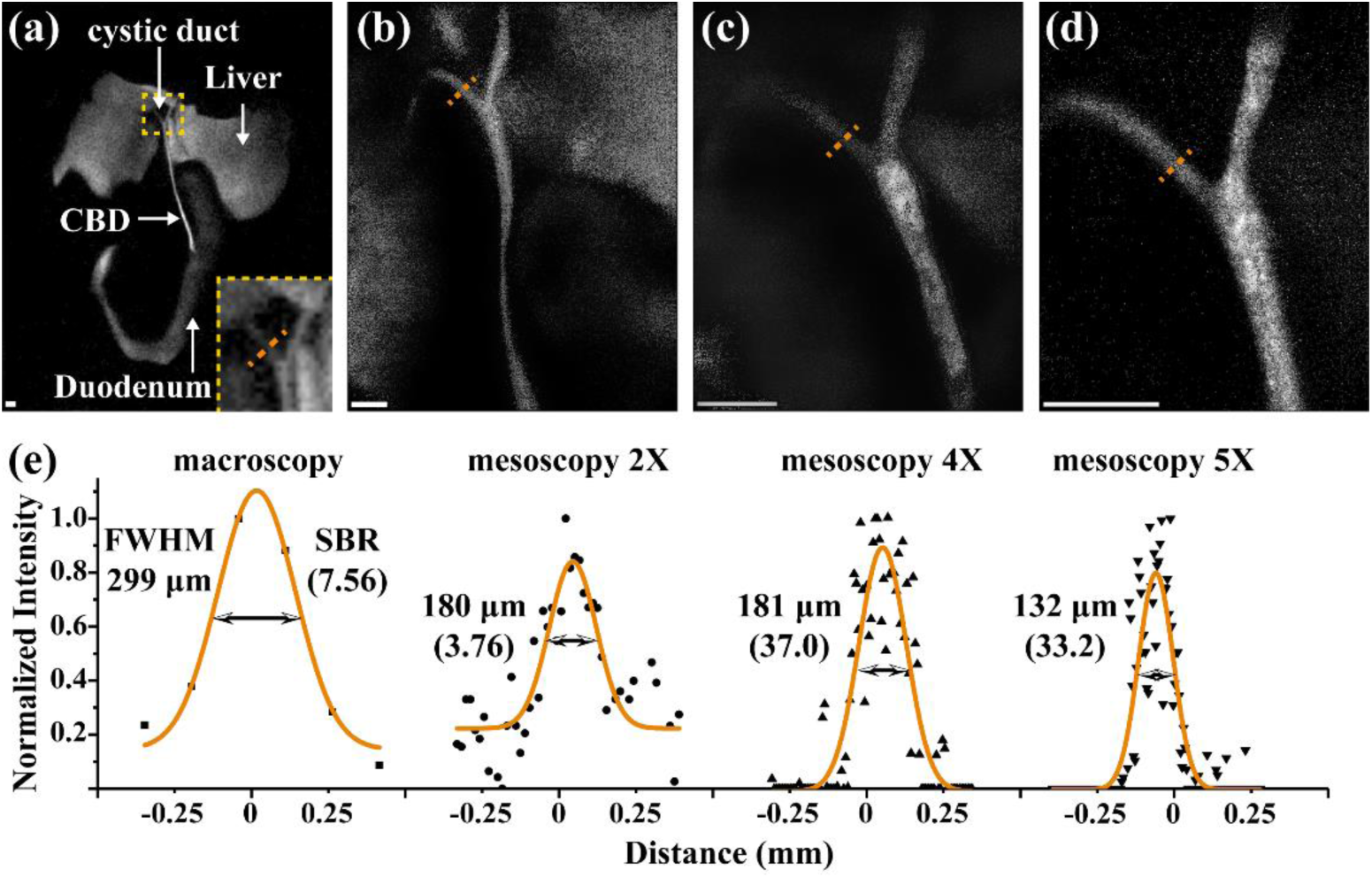
Real-time NIR-IIb fluorescence (a) macroscopic imaging and (b-d) mesoscopic imaging of biliary tract in the mouse intravenously injected with ICG (0.2 mg/mL, 200 μL). The magnifications of the objective were 2X, 4X, and 5X respectively. Scale bar= 1000 μm. (e) Gaussian fitting curves of cystic duct pointed in a-d. CBD: common bile duct.

Consequently, it would be available to detect the potential bile leakage or other biliary lesions more accurately benefiting from the bright fluorescence signal in the biliary tract and the weakened background signal from the liver tissue in the NIR-IIb window.

### 3.4. NIR-IIb fluorescence wide-field microscopic cerebrovascular imaging

2-photon fluorescence cerebrovascular imaging (exciting ICG with femtosecond pulses at 1617 nm) realized 2000 μm imaging depth ^[39]^. Moreover, microscopic cerebrovascular imaging assisted with ICG in the NIR-II region has been performed with large imaging depth (850 μm) and high resolution (10.71 μm) in mice with a cranial window ^[23]^. We then evaluated the feasibility of NIR-IIb fluorescence wide-field microscopic cerebrovascular imaging with ICG. ICG (2.5 mg/mL, 100 µL) was intravenously injected into mice with intact skull. NIR-IIb image with a large field (Fig.4c) displayed clearer blood vessels compared with NIR-II image (Fig.4a-b). This could be possibly explained that intact skull increased the scattering of short-wavelength NIR-II fluorescence. Gaussian fitting analysis of top right vessel showed that NIR-IIb fluorescence imaging has smallest FWHM and highest SBR, compared with images performed with 1000 nm LP and 13000 nm LP (Fig.4d). Furthermore, a small blood vessel at medium position displayed 7.87 µm spatial resolution, while vessel structures in 1000 nm LP and 1300 nm LP groups could hardly be resolved through background noise and became quite indistinguishable (Fig.4e).

**Fig. 4:**
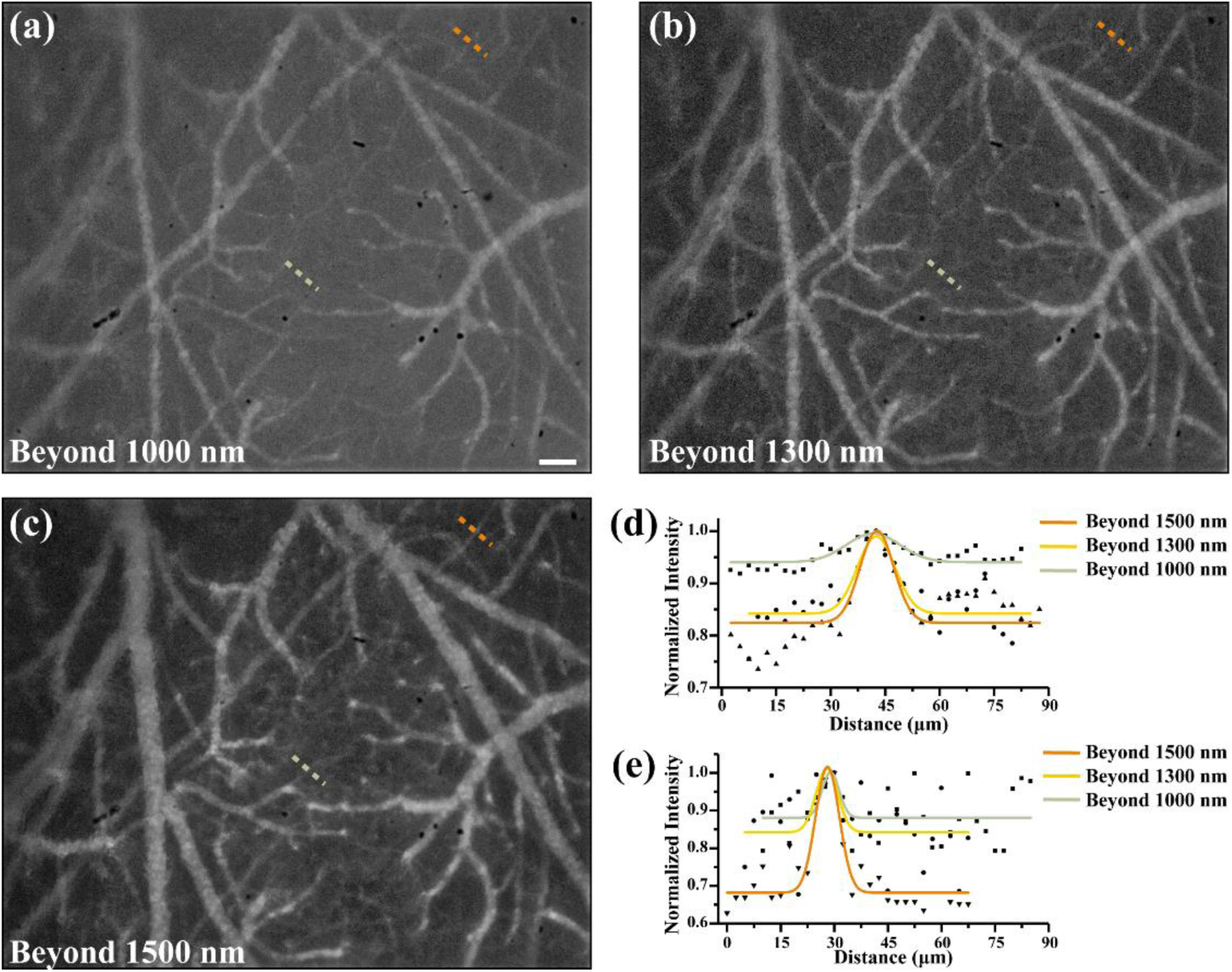
Real-time NIR-IIb fluorescence microscopic cerebrovascular imaging of the mouse with intact skull treated with ICG: (a-c) Images of brain blood vessels with 1000 nm, 1300 nm and 1500 nm long-pass filters respectively. The mouse was intravenously injected with ICG (2.5 mg/mL, 100 µL), excited by a 793 nm laser. Images were collected via an objective with the magnification of 5X. Scale bar= 100 μm. (d) Gaussian fitting curves of blood vessels in a-c at top right positions. FWHM= 10.90 µm (1500 nm LP), 12.11 µm (1300 nm LP), 17.44 µm (1000 nm LP); SBR= 1.21 (1500 nm LP), 1.17 (1300 nm LP), and 1.05 (1000 nm LP). (e) Gaussian fitting curves of blood vessels in a-c at medium positions; FWHM= 7.87 µm (1500 nm LP), 7.17 µm (1300 nm LP), and 5.61 µm (1000 nm LP); SBR= 1.49 (1500 nm LP), 1.20 (1300 nm LP), and 1.13 (1000 nm LP); Gaussian fitting R^2^= 0.830 (1500 nm LP), 0.230 (1300 nm LP), and 0.149 (1000 nm LP).

Considering the high photon scattering of skull, mice with a cranial window were also imaged. 400 μm imaging depth and capillary vessels as small as 3.75 μm were displayed (Fig.5a). Then, photo thrombotic ischemia (PTI) induction was carried out. A 532 nm laser beam was employed to excite photosensitizer (rose Bengal) for PTI induction. Released singlet oxygen through photosensitizer would adhere to the endothelial cells of blood vessels and damage them, causing cerebral thrombosis ^[40]^. Cerebral vessels were imaged before and after PTI formation (Fig.5b). Dashed lines manifested the corrupted vessels. Abnormal increases of fluorescence intensity in the vessels displayed the obstructed blood which was highlighted using solid line. Therefore, our study suggested the possibility of diagnosis of brain diseases via NIR-IIb fluorescence microscopic imaging with ICG.

**Fig. 5.**
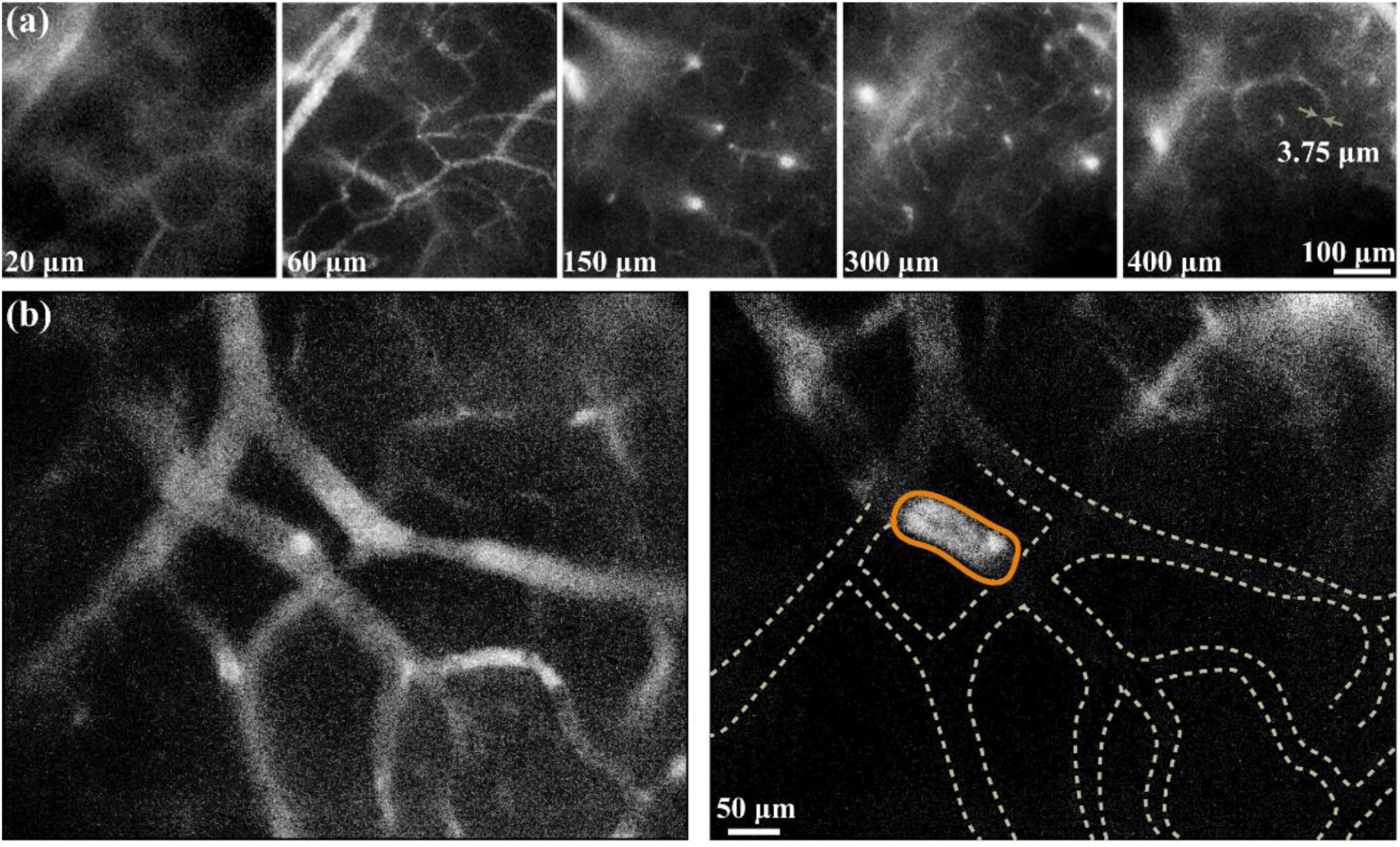
Real-time NIR-IIb fluorescence microscopic cerebrovascular imaging of the mouse with a cranial window intravenously injected with ICG (2.5 mg/mL, 100 μL): (a) Cerebral vasculatures images at various depths (20-400 μm) recorded by 25X objective (XLPLN25XWMP2, NA = 1.05, Olympus). (b) Cerebral vasculatures images before and after the formation of photo thrombotic ischemia (PTI).

## 4. Conclusion

In summary, we demonstrated the enhanced NIR-IIb fluorescence intensity of ICG when dissolved in the 4% HSA, rat bile and FBS solution. Benefiting from the reduced auto-fluorescence in the NIR-IIb region, the lymphatic drainage and the biliary tract could be imaged with higher contrast and resolution. Moreover, NIR-IIb fluorescence microscopic imaging assisted with ICG was firstly presented. Therefore, ICG bears great potential for translation of NIR-IIb imaging technology to clinical trial with the help of proteins. Fluorescence imaging in the NIR-IIb window could also be successfully performed using the clinical approved agent ICG and might be a promising technology for clinical diagnosis.

## Acknowledgments

This work was jointly supported by the National Natural Science Foundation of China (61735016 and 61975172), and the Zhejiang Provincial Natural Science Foundation of China (LR17F050001).

## Author contributions

Mubin He was mainly responsible for the design and operation of imaging experiments and data analysis. Di Wu mainly assisted with biliary imaging experiment, provided comments and revised the manuscript. Yuhuang Zhang assisted with data analysis and characterization experiment. Xiaoxiao Fan mainly assisted with lymph imaging experiment. Jun Qian was the corresponding author of this article, designed the imaging systems and provided guidance in the entire research process.

## Appendix A. Supplementary materials

Supplementary materials to this article can be found online.

